# A large replicase of nidovirus-like complexity in a putative RNA virus order *“Quisvirales”* expands the known diversity of helicase superfamilies

**DOI:** 10.64898/2026.01.28.702245

**Authors:** Alexander E. Gorbalenya, Dmitry V. Samborskiy, Chris Lauber

## Abstract

Helicases are essential ATPases that unwind nucleic acids and are classified into six recognized superfamilies (SF1–SF6). All positive-stranded RNA viruses (ssRNA+) with genomes larger than ∼7 kb encode SF1-SF3 helicases, linking them to RNA genome expansion. In the phylum *Pisuviricota* that includes many pathogens, helicases of SF1-SF3 are integrated into multi-enzyme replicases including 3C(-like) proteases (3CLpro) and RNA-dependent RNA polymerases (RdRp). Here, large-scale mining of invertebrate metastranscriptomes uncovered six related spider-associated ssRNA+ viruses that, based on their conserved 3CLpro–RdRp module, genome size (20–22 kb), and phylogeny, form a putative new order, *“Quisvirales”*. Quisviruses have similar genome and replicase architectures to enveloped coronaviruses and other nidoviruses. However, the nidovirus SF1 helicase is replaced in quisviruses by a novel superfamily helicase that, like the *Picornavirales* SF3 helicase, comprises an AAA+ (ATPase-like) domain. Thus, viruses with large replicases of nidovirus-like complexity may have evolved repeatedly from an 3CLpro–RdRp-encoding ancestor.

## Introduction

The RNA-dependent RNA polymerase (RdRp) is the hallmark protein of RNA viruses and the principal nucleic acid-processing enzyme required for viral genome replication. In all known positive-stranded RNA viruses (ssRNA+) with documented proteome and a genome size of more than 7.1 kb and some with smaller genomes starting with 5.4 kb ^1^, the RdRp is supported by a cognate essential helicase as part of the viral replicase complex ^2–5^. Helicases may be indispensable for replication and expression of (relatively) large RNA genomes and are among most common enzymes of RNA viruses ^6^. They are ATP-hydrolyzing motor proteins (ATPases) that move directionally along single- or double-stranded DNA or RNA molecules while displacing a single-stranded polynucleotide or protein from multi-subunit complexes. Universal features of all known helicases include three conserved sequence motifs, two involved in binding and hydrolysis of NTP (Walker A and B boxes) and one containing an arginine finger for energy coupling to the helicase displacing activity. Helicases may act on DNA, RNA, or DNA-RNA hybrid templates, move from 5’-to 3’-end or reverse direction and have been associated with a vast variety of functions in (infected) cells.

Based on sequence and structural similarities and mechanistic differences, helicases have been classified into six superfamilies (SF1 to SF6) present in DNA-based life forms, three of which (SF1, SF2, and SF3) are also found in ssRNA+ viruses ^7,8^. The core of SF1 and SF2 helicases is composed of two paralogous subdomains with RecA-like folds encompassing seven to eight sequence motifs ^9^. The active form of SF1/SF2 helicases is the monomer, although they may also form dimers ^10–13^. Translocation of SF1/SF2 helicases along the DNA/RNA template is assumed to follow an inchworm-like mechanism ^14^. In contrast, the core of SF3-6 helicases is twice smaller and comprises a single P-loop NTPase domain, either of RecA-like fold (SF4-5) or of related AAA+ domain (ATPases associated with diverse cellular activities plus) in SF3 and SF6 ^9,15,16^. It includes four to six sequence motifs, four of which are the universal Walker A and B boxes, a sensor-1 and an Arg finger, with SF-specific spacing. Helicase molecules of SF3-6 assemble into a multimeric ring that may undergo oligomeric transition and is the active enzymatic form ^8,17^. Translocation of SF3-6 helicases is proposed to follow a hand-over-hand mechanism in which sequential ATP hydrolysis by subunits causes ring rotation coupled with its movement along the template ^16^.

Helicase-encoding RNA viruses belong to several orders and three families of phyla *Kitrinoviricota* and *Pisuviricota* (kingdom *Orthornavirae)* ^18^. In the *Pisuviricota*, the proteome of orders *Nidovirales* and *Picornavirales* (class *Pisoniviricetes*) and *Patatavirales* (class *Stelpaviricetes*) include helicases of SF1, SF3 and SF2, respectively, either upstream or downstream of the essential 3C(-like) protease (3CLpro) and protein-primer-dependent RdRp; the observed patterns are order-specific. 3CLpro controls genome expression by mediating autoproteolytic processing of polyprotein(s) into subunits of the replication-transcription complex; in viruses that use jelly-roll capsids, it also cleaves precursors of capsid subunits ^19^. There are two other 3CLpro-RdRp-encoding ssRNA+ orders, *Sobelivirales* (class *Pisoniviricetes*) and *Stellavirales* (class *Stelpaviricetes*), that have genome sizes under 7.1 kb and no helicase encoded. The phylum *Pisuviricota* includes main pathogens of humans, e.g. SARS-CoV-2, poliovirus and norovirus.

Described some 35 years ago, helicase classification and domain linkages still hold in many thousands of the newly discovered viruses from an ever-expanding host range of mono- and multi-cellular eukaryotes ^3,6,20–22^. They imply that the observed genomic and phylogenetic patterns must have emerged early in the evolution of ssRNA+ viruses. Also, they reveal constraints on main variation of 3CLpro-RdRp-encoding viruses, especially in relation to helicases. While the 3CLpro-RdRp-encoding viruses account for a subset of RNA viruses, they cover a nearly full range of the documented RNA genome sizes from 4 to 64 kb and have explored a variety of genomic organizations and expression mechanisms during evolution in mono- and multi-cellular eukaryotes ^23–31^.

Here, we delineate a putative ssRNA+ virus order *“*Quisvirales*”* in the class *Pisoniviricetes*, based on RdRp plylogeny and the presence of 3CLpro-RdRp loci in genome sequences reconstructed from spider metatranscriptomes. Quisviruses have large RNA genomes in the size range of 20-22 kb and a putative multi-enzyme replicase of large nidovirus-like complexity. However and contrary to the observed domain linkages in relation to genome size, quisviruses do not encode a helicase of known superfamilies. Instead we identified a novel AAA+ domain that is conserved in six known quisviruses in the genomic position of nidovirus SF1 helicase. Like ring-forming helicases of SF3-6, this putative ATPase is predicted to adopt a single RecA-like fold at its core and includes four key sequence motifs. Its evolutionary distances from SF1-6 helicases are as large as between different SFs. We propose that this ATPase is the prototype of a new helicase superfamily and may play a key role during replication of quisviruses.

## Methods

### Virus discovery from public sequence databases

We applied a data-driven virus discovery approach using the Virushunter and Virusgatherer workflows to screen 30,105 arthropod transcriptome sequencing projects from the Sequence Read Archive (SRA) for the presence of RNA virus genomes, as described previously ^23,32–34^. Among the hits were two divergent viral genomes that clustered together and otherwise had highest sequence similarity to members of the order *Picornavirales* and other pisoniviruses in a protein BLAST search (blastx) against the National Center for Biotechnology Information (NCBI) viral_protein database. They constitute the first two representatives of quisviruses, recognized and named in this study (see below). We used the predicted RdRp-encoding polyproteins of these two viral genomes as query in a protein-level Blast search (tblastn with default parameters), conducted at the NCBI website, against all Arthropoda sequences from the Transcriptome Shotgun Assembly (TSA) database ^35^. We considered hits with an E-value of 1×10^-50^ or better and a contig length of at least 10 kb. This TSA screening approach yielded four additional quisvirus hits. These four viral genomes were re-assembled from the corresponding raw-read SRA data using Megahit with default parameters ^36^. We obtained several additional strongly supported hits against contigs shorter than 10 kb in different spider projects, which were not considered for further analysis. Details about the six viral genomes and the datasets in which they were discovered can be found in **Table S1**.

### Viral genome organization

Open reading frames were predicted using getorf from the EMBOSS package ^37^, considering ORFs from start to stop codons of at least 300 nucleotides, except for ORF1b that was delimited from stop to stop codons. Programmed ribosomal frameshifting elements were predicted using KnotInFrame ^38^.

### Protein domain annotation

Quisvirus protein domain borders were delineated using multiple sequence alignment to reference RNA genomes and their annotated proteins in the Viralis software ^39^. Functional annotation of quisvirus proteins involved profile HMM comparisons to viral and host reference proteins from the PDB, Pfam-A, Uniprot and NCBI Conserved Domains databases utilizing HHpred searches at the MPI Bioinformatics Toolkit web service (https://toolkit.tuebingen.mpg.de) ^40–44^ or as described in our research ^23^.

### Phylogenetic analysis

We included the discovered quisviruses and one representative per ICTV-recognized virus family of the orders *Sobelivirales* (3 virus families), *Picornavirales* (9), *and Nidovirales* (14) from the class *Pisoniviricetes*, a divergent unclassified picorna-like virus as well as genus representatives of the orders *Patatavirales* (3 virus genera from the family *Potyviridae*) and *Stellavirales* (2 from the family *Astroviridae*) from the class *Stelpaviricetes* into the phylogenetic analysis (**Table S2**). We constructed a multiple sequence alignment of the RdRp amino acid sequences demarcated by motifs G and E ^45,46^ at, respectively, the N- and C-terminus within the Viralis software^39^. From this alignment, we extracted all columns (n=208) that contained less than 10% gaps for phylogenetic analysis. The best fitting evolutionary model was selected using ModelTest-NG version 0.2.0 ^47^, which was the BLOSUM62+I+G4 model. A maximum likelihood phylogenetic tree of the RdRp proteins was reconstructed using PhyML version 20120412 with parameters *‘*-d aa -m BLOSUM62 -f m -v e -a e -c 4 -o tlr’ ^48^. Branching support was assessed using the Shimodaira–Hasegawa (SH)–like procedure.

### Helicase similarity network

Multiple sequence alignments of helicase superfamilies SF1-6 were obtained from either Pfam/InterPro (SF6 RuvB: accession PF05496), CDD (SF4 Gp4D: accession CD19483, SF5 Rho: accession CD01128), PROSITE (SF3 of DNA viruses: accession PS51206) ^41,43,49^, or in-house from Viralis (SF1, SF2, SF3 of RNA viruses). For SF2 we included helicases from *Potyviridae and Flaviviridae* and flavi-like viruses. For SF3 we generated two alignments encompassing proteins from (i) *Caliciviridae* and calici-like viruses and (ii) *Picornaviridae* and picorna-like viruses. Scores obtained from the comparison of the two SF3 multiple sequence alignments serve as a positive control.

The multiple sequence alignments for SF1-6 and the quisvirus ATPase alignment were subjected to pairwise HMM-vs-HMM comparisons using hhalign in (default) local mode with parameter *‘*-M first’ from the HHblits package ^50^. For each pair of alignments, we considered two comparisons treating an alignment either as query or as template in the hhalign analysis. From these two comparisons we collected the Sum_probs values, which integrate over the posterior probabilities of all aligned pairs of match states. To obtain a final score for the alignment pair, we took the maximum of these two Sum_probs values (maximum sum of poster probabilities, MSPP). The MSPP values were scaled by multiplying them with a factor of 0.025 for all alignment pairs (**Table S3**) and used as edge weights in a similarity network of the helicases, which was visualized in R using the igraph package ^51^.

### Protein 3D structure modeling

Protein tertiary structures were predicted using AlphaFold3 Server ^52^. We conducted two independent modeling runs for each input sequence and considered the five top-ranking structures. The in total 10 structures were aligned and visualized using ChimeraX to assess prediction variability ^53^. We aligned the (putative) ATPase structures of Strandella quadrimaculata quisvirus (accession IAMJ01023800) and representatives for SF3 (2C of poliovirus 1; NP_041277), SF4 (Gp4D of Prochlorococcus phage P-RSP2; AGF91554), SF5 (transcription termination factor Rho Chlorobaculum tepidum TLS; NP_661168), and SF6 (Holliday junction branch migration DNA helicase RuvB of Bdellovibrio bacteriovorus; WP_011164893) using Kpax with the -multi option ^54^. This structural alignment was used by Kpax to generate a multiple sequence alignment for these five ATPases, which was manually refined around globally conserved residues. The multiple sequence alignment was visualized using Jalview ^55^.

## Results

### Discovery of a new lineage in the phylum Pisuviricota

From a Data-Driven Virus Discovery (DDVD) ^23,34,56^ screen for RNA viruses in 30,105 arthropod transcriptome sequencing projects from the SRA, we initially identified two novel and closely related viral genomes. They diverged profoundly from all known RNA viruses and showed only remote RdRp sequence similarity to members of the order *Picornavirales*. The two viral genomes were found in spider sequencing projects of *Nephilengys cruentata* and *Pardosa pseudoannulata* (**Table S1**). We used the predicted RdRp-encoding proteins of these two viral genomes as query in a Blast search against the TSA database and identified four additional viral genomes of this virus group in data from the 1000 spider silkomes ^57^ projects of *Strandella quadrimaculata, Clubiona zilla, Octonoba sybotides*, and *Haplodrassus kanenoi*. We only obtained hits against spider projects and no hits in other arthropods.

### Genome architecture and protein annotation of new viruses

Based on the presence of terminal non-coding sequences, 5’-UTR and 3’-UTR, and other features (see below), we considered four of the six novel viral genomes to be coding-complete with genome sizes ranging from 20474 to 22035 nt, while two are partial (12005, 14810 nt) (**Table S1**). The complete genomes have a multi-ORF organization including at least five ORFs flanked by 5’- and 3’-UTRs. The two largest and overlapping ORFs, ORF1a and ORF1b with a −1 programmed ribosomal frameshifting (−1 PRF) element with slippery sequence AAAAAAC or UUUAAAC in the overlap (**Supplementary Figure 1**), account for ∼80% of the genome size. They are followed by four to six partly overlapping predicted ORFs of sizes in the range of 101 to 661 aa (**Table S4**). Translation of ORF1a and ORF1b from the genomic RNA may produce two polyproteins, termed pp1a (from ORF1a) and pp1ab (ORF1a and ORF1b), like in nidoviruses.

We observed a sharp and step-wise increase of read coverage depth at the 3’ genome region for five of the six viruses with the highest mean coverage (**Supplementary Figure 2**). This sequencing coverage depth pattern is highly similar to that seen for nidoviruses ^23,34^ and suggests that the newly discovered viruses express the ORFs downstream of ORF1b in the 3*’* genome region via subgenomic RNAs.

We used amino acid multiple sequence alignments of the six viruses to scan different sequence and structural databases in profile mode for genome annotation (see M&M). Based on significant hits (E-value>10^-3^), we identified six putative non-structural protein domains that are conserved across the six viruses (**Table S5**). They include papain-like protease (PLpro) and 3CLpro in pp1a as well as four domains downstream of −1PRF in the pp1ab. They are putative RdRp, RNA genome expansion and replicase associated (REXA) ^23^, novel AAA+ ATPase, and DNA/RNA demethylase AlkB-like domain ^58^ (**Supplementary Figures 3 and 4**). A potential trans-membrane (TM) domain has been identified upstream of 3CLpro (**Table S5**) in the apparent resemblance between the two-domain organization of TM-3CLpro of quisviruses and 2B-3Cpro of picornaviruses. Somewhat similar, top hits for five enzymes PLpro, 3CLpro, RdRp, REXA and AlkB were predominantly observed against ssRNA+ virus proteins, mostly from the phylum *Pisuviricota*, and specifically family *Picornaviridae*. Hit aa conservation varied greatly, from very high and domain-wide (up to e-value 10^-13^ and 90% coverage, AlkB) to moderate (RdRp and REXA) to limited around the active center (PLpro and 3CLpro) (**Table S5**). In all five enzymes, known essential residues are conserved indicating that they are functional. We describe results for the novel AAA+ ATPase in a separate section below.

Based on characterization of other viruses, we propose that pp1a/pp1ab are autoproteolytically processed to mature proteins by 3CLpro with possible assistance of PLpro. The cleavage sites in six viruses may be conserved but remain unknown. Accordingly, the exact boundaries/sizes and identity of mature proteins of these viruses are yet to be established.

In addition to the non-structural domains, we identified two putative structural domains of presumed virus particles in the ORFs downstream of ORF1b that are conserved in all six quisviruses. This includes two glycoprotein (GP)-like domains in, respectively, the C-terminal part of ORF3 (GP1) and in ORF4 (GP2) (**Table S5**). Both domains include a putative TM region at the C-terminus and eight and ten conserved Cys residues, respectively; these features are common for GPs of enveloped RNA virus particles (**Supplementary Figure S5**). Moreover, the N-terminal part of ORF3 encodes an Immunoglobulin-like domain that is conserved in four quisviruses which are relatively closely related to each other but not in Octonoba sybotides quisvirus and Pardosa pseudoannulata quisvirus that form two basal lineages in the RdRp phylogeny (**Supplementary Figure 5, Figure 2**). Most quisvirus genomes have at least one short ORF downstream of ORF4 (**Figure 1**) that may encode capsid proteins, although we did not observe significant sequence similarity to reference database entries. These results suggest that quisviruses employ enveloped virions, like nidoviruses and unlike picornaviruses, and in line with the plausible sgRNA expression of the involved ORFs.

**Figure 1.**
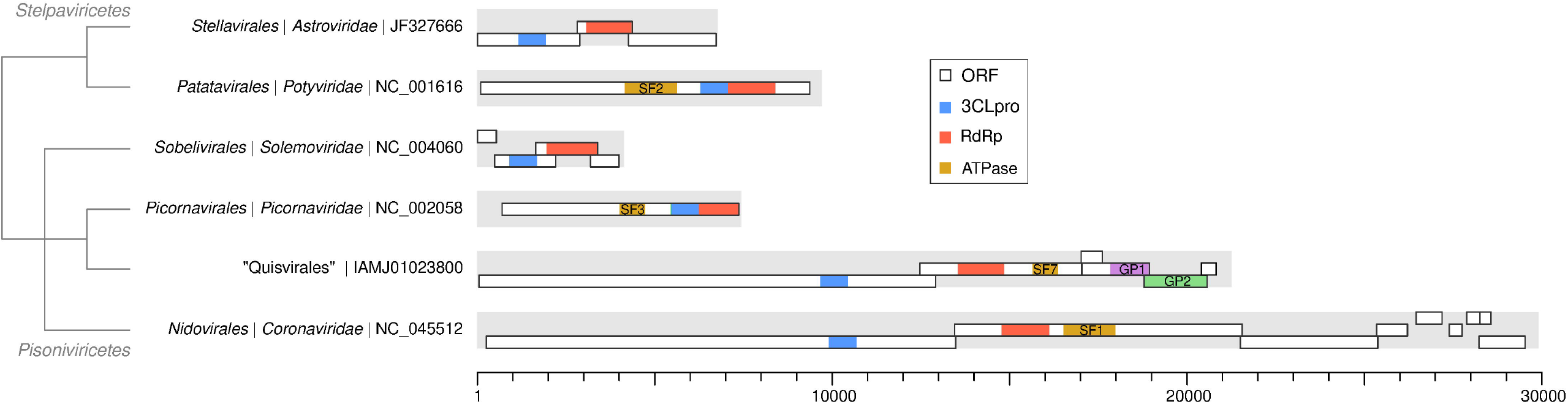
Genomic organization of quisvirus and selected *Pisuviricota* representatives. Shown are genome maps for a quisvirus and one representative for each virus order whose members encode a 3C-like protease (blue) and an RdRp (red) domain. Some of the viruses also encode a helicase/ATPase domain (yellow). For the Strandella quadrimaculata quisvirus representing *“Quisvirales”*, ORFs of at least 150 nt in length are shown and two predicted glycoprotein (GP)-like domains, GP1-like (violet) and GP2-like (green), are highlighted. Genome and ORF sizes are drawn to scale. Borders of protein domains are estimates based on either annotation in Genbank records or hits of profile HMM-based sequence homology searches. The cladogram at the left is based on the classification of the included orders in the ICTV taxonomy and the phylogenetic analysis in this study (**Figure 2**); branch lengths have no meaning.

**Figure 2.**
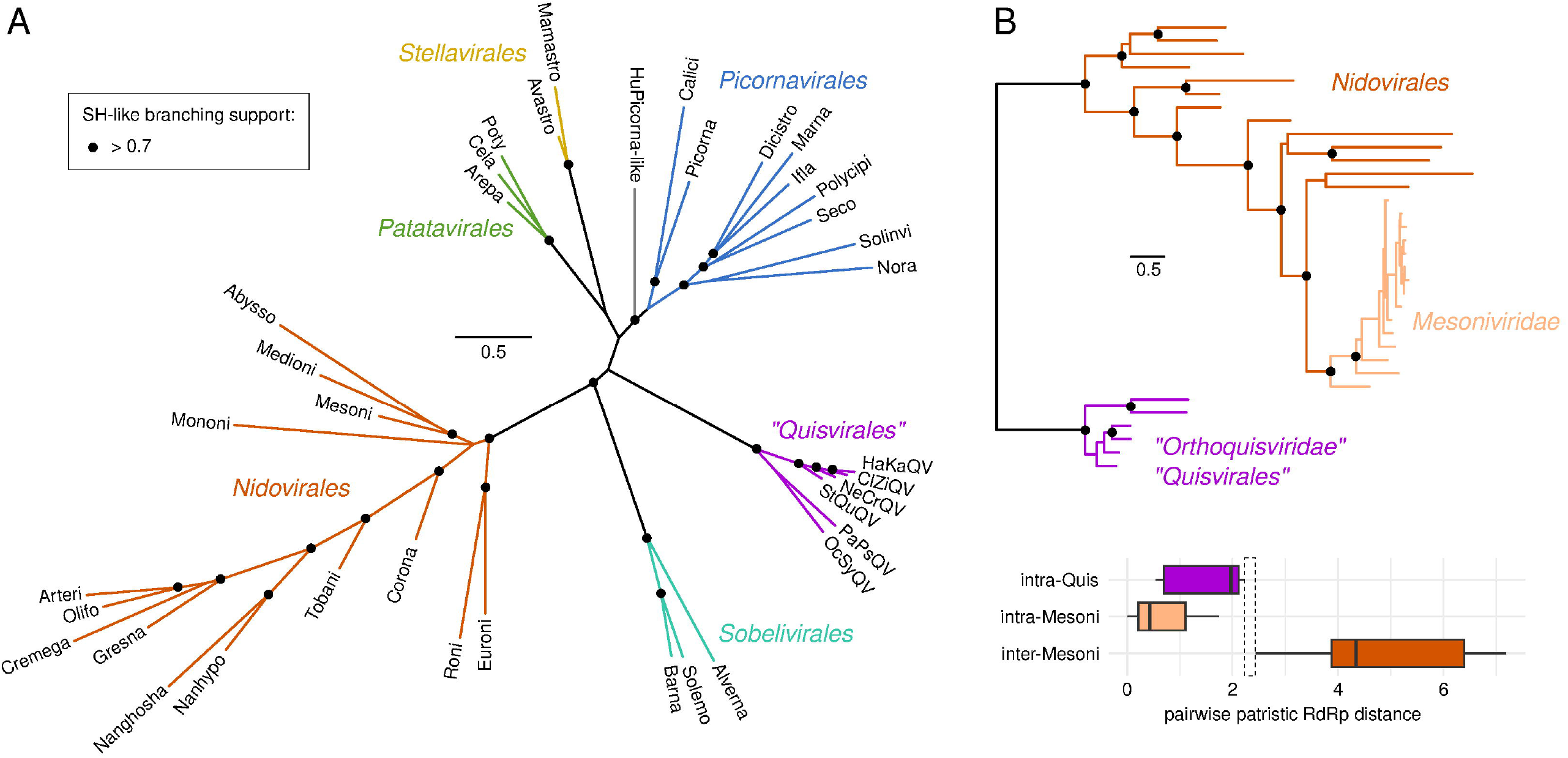
Phylogenetic relationships of quisviruses with other members of the phylum Pisuviricota encoding the 3CLpro-RdRp domain pair. The maximum likelihood RdRp-based phylogeny was reconstructed using PhyML. Quisviruses as well as ICTV-recognized orders *Picornavirales, Nidovirales, Sobelivirales, Patatavirales* and *Stellavirales*, as well as an unclassified virus (gray), are discriminated by colored branches. One representative virus per ICTV-recognized virus family or genus was included in the analysis (**Table S2**). Dots at internal branching events indicate SH-like support values larger 0.7; otherwise no dot is shown. The scale bar is in average number of amino acid substitutions per site. The included quisviruses are: Clubiona zilla quisvirus (ClZiQV; accession: AWI01044675), Haplodrassus kanenoi quisvirus (HaKaQV; IBAC01001151), Nephilengys cruentata quisvirus (NeCrQV; SRR3943478), Octonoba sybotides quisvirus (OcSyQV; IAZZ01033660), Pardosa pseudoannulata quisvirus (PaPsQV; SRR22498918), Strandella quadrimaculata quisvirus (StQuQV; IAMJ01023800). For ICTV virus orders, one representative per reference virus family or genus is shown: *Sobelivirales*: *Solemoviridae* (Solemo; Southern bean mosaic virus; NC_004060.2), *Barnaviridae* (Barna; Mushroom bacilliform virus; NC_001633.1), *Alvernaviridae* (Alverna; Heterocapsa circularisquama RNA virus; NC_007518.1); *Picornavirales*: *Noraviridae* (Nora; Nora virus; NC_007919.3), *Polycipiviridae* (Polycipi; Lasius neglectus virus; NC_035450.1), *Secoviridae* (Seco; Camellia virus A; NC_077103.1), *Solinviviridae* (Solinvi; Nylanderia fulva virus 1; NC_030651.1), *Picornaviridae* (Picorna; Poliovirus; NC_002058.3), *Dicistroviridae* (Dicistro; Cricket paralysis virus; NC_003924.1), *Iflaviridae* (Ifla; Deformed wing virus; NC_004830.2), *Marnaviridae* (Marna; Heterosigma akashiwo RNA virus; NC_005281.1), *Caliciviridae* (Calici; Norwalk virus; NC_001959.2); *Nidovirales*: *Tobaniviridae* (Tobani; Breda virus; NC_007447.1), *Coronaviridae* (Corona; Severe acute respiratory syndrome coronavirus 2; NC_045512.2), *Abyssoviridae* (Abysso; Aplysia californica nido-like virus; NC_040711.1), *Mesoniviridae* (Mesoni; Nam Dinh virus; NC_015874.1), *Medioniviridae* (Medioni; Botrylloides leachii nidovirus; NC_055538.1), *Mononiviridae* (Mononi; Planarian secretory cell nidovirus; NC_040361.1), *Euroniviridae* (Euroni; Beihai hermit crab virus 4; NC_032490.1); *Roniviridae* (Roni; Yellow head virus; NC_043505.1), *Gresnaviridae* (Gresna; Guangdong greater green snake gresnavirus; NC_046959.1), *Cremegaviridae* (Cremega; Trionyx sinensis hemorrhagic syndrome cremegavirus; NC_076637.1), *Arteriviridae* (Arteri; Lactate dehydrogenase-elevating virus; NC_001639.1), *Olifoviridae* (Olifo; Hainan oligodon formosanus olifovirus; NC_046958.1), *Nanghoshaviridae* (Nanghosha; Nanhai ghost shark nanghoshavirus; NC_046960.1); *Nanhypoviridae* (Nanhypo; Wuhan japanese halfbeak nanhypovirus; NC_046957.1); *Potyvirus* (Poty; Potato virus Y; U09509); *Celavirus* (Cela; Celery latent virus; MH932227), *Arepavirus* (Arepa; Areca palm necrotic spindle spot virus; MH330686); *Mamastrovirus* (Mamastro; Human astrovirus type 1; Z25771); *Avastrovirus* (Avastro; Avastrovirus 1; Y15936); unclassified (HuPicorna-like; Hubei picorna-like virus 69; NC_033021).

### A new order “Quisvirales”of the class Pisoniviricetes

The observed 3CLpro-RdRp domain segregation was previously described in three orders of the class *Pisoniviricetes* and two orders of the class *Stelpaviricetes* (all in the phylum *Pisuviricota*) (**Figure 1**). When tested in the RdRp-based phylogeny of the class *Pisoniviricetes*, which included a prototype virus of each recognized virus family in the class, the six novel viruses formed a divergent, monophyletic lineage positioned somewhat closer to Picornavirales, compared to the other two orders *Sobelivirales* and *Nidovirales* (Figure 2A)

Based on the RdRp phylogeny, we propose that these viruses form a new RNA virus order in the class *Pisoniviricetes*. Due to several highly unusual features of the novel viruses, we provisionally named this novel order *“Quisvirales”*, referring to the Latin word *“*Quis*”*, which translates to *“*What is this?*”*. The pairwise evolutionary distances in the RdRp between the six quisiviruses is at a scale observed for the family *Mesoniviridae* (order *Nidovirales*) that have viruses with similar genome sizes (20.0-22.0 kb) (**Figure 2B**). We therefore propose that the described quisviruses form a provisional virus family which we named *“Orthoquisviridae”*. The six quisviruses may be prototypes of six different virus species according to their genetic divergence (**Figure 2B**).

### Quisviruses may encode a novel type of ATPases that resembles single AAA+/RecA-like helicases

Based on a strong link between the presence of a cognate helicase in ssRNA+ viruses with large genome size (> 7.1kb) and the similarities in genomic organization to nidoviruses, we expected to identify a SF1 helicase domain in pp1ab of quisviruses in our sequence-vs-profile searches. Surprisingly, we observed no hits against SF1-SF2 profiles and only moderately supported hits against SF3-SF4 profiles that were limited to Walker box A and B and which are common for a wide range of ATPases (**Table S6**). This analysis identified a new ATPase (∼190aa) downstream of and close to the RdRp, the position that is occupied by a much larger SF1 helicase (HEL1) in nidoviruses (**Figures 1 and 3**).

**Figure 3.**
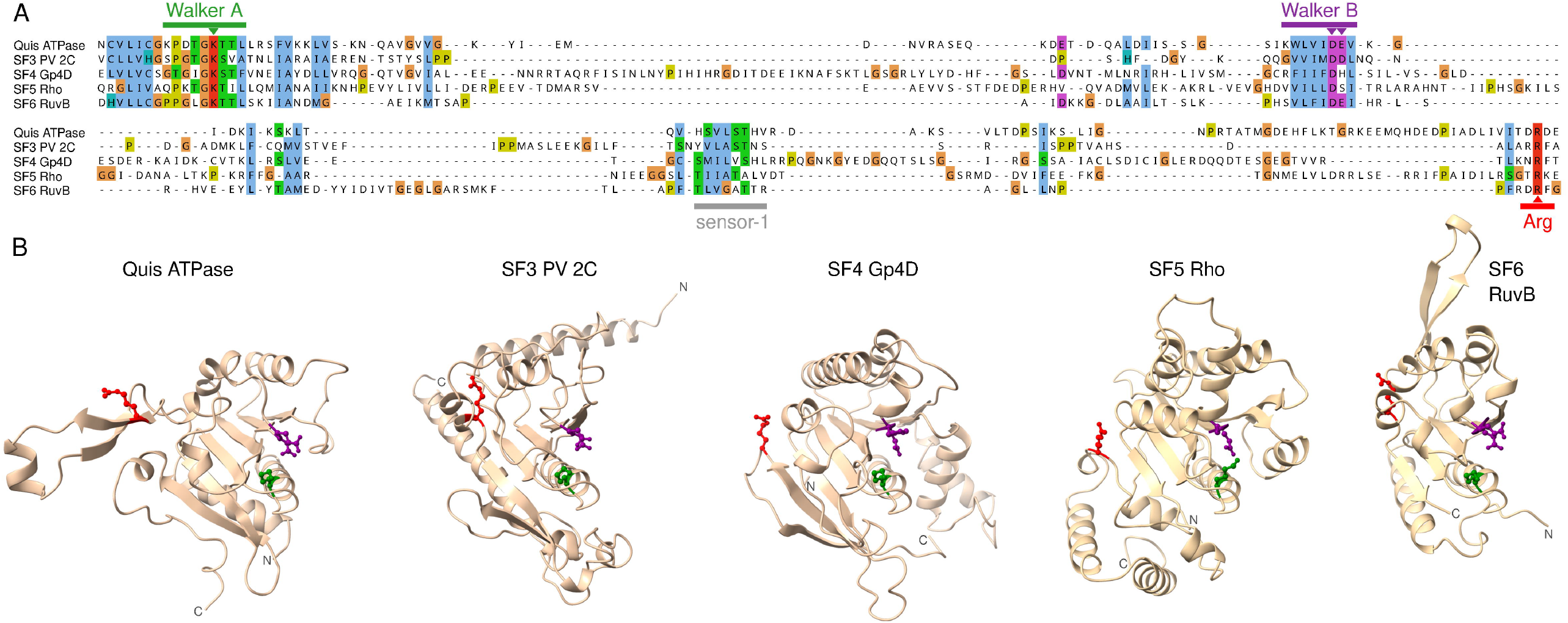
Sequence and structural features of the quisvirus ATPase are indicative of RecA-like structure and helicase activity. (A) Multiple sequence alignment of quisvirus ATPase (IAMJ01023800) and superfamily 3 2C helicase of poliovirus 1 (SF3 PV 2C; Genbank accession NP_041277), superfamily 4 Gp4D helicase of Prochlorococcus phage P-RSP2 (SF4 Gp4D; AGF91554), superfamily 5 transcription termination factor Rho of Chlorobaculum tepidum TLS (SF5 Rho; NP_661168), and superfamily 6 Holliday junction branch migration DNA helicase RuvB of Bdellovibrio bacteriovorus (SF6 RuvB; WP_011164893). The sequence alignment was generated using Kpax and is based on an alignment of predicted tertiary structures of the proteins. The N- and C-terminal parts of the alignment are not shown for simplicity. The alignment was visualized using Jalview with Clustal coloring mode. (B) The predicted structures of quisvirus ATPase and SF3-6 helicases were superimposed and are shown side-by-side. Four conserved residues in three of four motifs indicated by rectangles in panel A are highlighted by color; green: K in Walker A; purple: DE in Walker B; red: R in arginine finger; gray: sensor-1.

We annotated this ATPase based on hits to 114 PFAM entries and many hundreds entries in other databases with hit E-values from 10^-3^ to 10^-11^ (**Table S7**). While the majority of hits were limited to the Walker A NTPase motif, there were significant hits against AAA+ domains in several bacterial uncharacterized multi-domain proteins that encompassed >75% of query and target domains (UniRef100, E-value from 10^-9^ to 10^-11^; the longest hit is 173aa to A0A022Q4Y0, 2.10^-10^). AAA+ domains facilitate many functions, including helicases of SF3-6 and those unclassified with single AAA+/RecA-like fold. Accordingly, longest hits for this ATPase were of 170 aa with SF4 DNA B helicases (PF03796, E-value 3.10^-7^) and 163 aa with SF4 helicase domains of E.coli phage T7 DNA PRIMASE/HELICASE (pdb70 entry 1CR0_A, E-value 2.10^-6^) and Twinkle ^59^ mtDNA replisome (8E2L_B, E-value 1.10^-4^) (**Table S7**). Hit alignments between the query and targets (templates) included four SF4 motifs of the template (**Figure 3A**).

Also, the quisvirus ATPase includes conserved sequence motifs (Walker A, Walker B, sensor-1, arginine finger) that are involved in ATP binding and hydrolysis as well as energy coupling to unwinding in known helicases (**Figure 3A**). Upon structural comparison of SF3-SF6 helicases and a quisvirus ATPase tertiary structural model (**Supplementary Figure 6A**), key residues of these sequence motifs occupied spatially highly similar positions in a single AAA+/RecA-like domain, implying functional similarity (**Figure 3B**). Accordingly, the quisvirus ATPase exhibits highest (although moderate) sequence affinity to members of the ring-forming single-AAA+/RecA-like SF3-SF4 in a network analysis (**Table S3**) in which the weights of connecting edges are proportional to MSPP scores obtained in pairwise profile HMM-based sequence comparisons (**Supplementary Figure 6B**). Namely, the MSPP score for quisvirus ATPase was in the range of 25.6-85.6 (mean MSSP of 46.9) against other SFs, comparable to that between DNA- and RNA-virus SF3 domains (67.4-70.3) and also to inter-SF MSPP scores (**Table S3**). Notably, the MSPP scores involving the quisvirus ATPase were considerably lower than that of the positive control (126.6) of comparing SF3 domains of calici-like and picorna-like viruses (**Supplementary Figure 6B**). Combined, these results suggest that the quisvirus ATPase is the prototype of a new helicase superfamily (SF7) and has helicase activity.

## Discussion

In this study we introduce a provisional order *“Quisvirales”* comprising six newly discovered RNA viruses of spiders which encode replicases of nidovirus-like complexity and deviate from all known ssRNA+ viruses with comparably large genome sizes in respect to the omnipresent helicase. Quisviruses either don’t encode a helicase domain, or encode a helicase different from known SF1-SF6 helicases of all life forms (favored by our analysis). Both the helicase/RNA-genome size linkage and the helicase superfamily classification were proposed several decades ago ^2,7^. Ever since they have sustained the fast pace of genomic and helicase characterization in numerous cellular organisms and viruses. Thus and regardless of which of two hypotheses proves correct, the discovered quisviruses already revise the established relationship between RNA viruses and helicases.

We acknowledge that further biochemical and structural characterization of the quisvirus proteome is essential to resolve the residual uncertainty. However, demonstration of helicase activity may take years of work as researchers have experienced with SF3 helicases of picornaviruses, most relevant to this study ^60,61^. The lack of helicase activity in experiments in the meantime may be due to technical reasons, and this is in contrast to the successful demonstration of ATPase activity and ring structure, two other genomic-based inferences. Due to these considerations, we did not pursue experimental research in this study. Instead we discuss below how the presented findings already advance our understanding of ssRNA+ viruses and why quisviruses encoding a new SF helicase seems most plausible.

To start with, we reiterate that our results are reliable, consistent with literature, original, and informative, even if technically they are limited to various bioinformatics analyses. We applied a DDVD approach ^56^ to initiate this study and observed statistical support for two assembled virus genomes in respect to RNA read depth and coverage breadth as derived from (infected) spider transcriptomes (**Supplementary Figure 2**). These assembly metrics are comparable to those reported in our other high-throughput virus genomic discoveries ^23,34^. Furthermore, four other closely related genome sequences were identified in the TSA and reassembled. The six new viruses share a multi-ORF genomic organization and encode several protein domains with characteristic sequence motifs typical for ssRNA+ viruses.

Due to the deep separation of this new group of viruses from the most closely related viruses in a RdRp-based tree and unique but conserved features of genome architecture, we propose a new tentative order provisionally named *“Quisvirales”*, class *Pisoniviricetes*, phylum *Pisuviricota*. In contrast to *“Quisvirales”* and *Nidovirales*, four other orders of the 3CLpro-RdRp encoding viruses (*Picornavirales, Sobelivirales, Stellavirales, Patatavirales*) have much smaller genomes and use naked capsids ^24–26,62^. The 20-22 kb genome sizes of quisviruses are exceeded only by flavi-like viruses and nidoviruses, while they resemble those of the nidovirus family *Mesoniviridae* ^63–65^. The observed quisvirus genome size range is narrow for a virus order and may reflect a currently limited sampling, virus family-wise (*“Orthoquisviridae”*) and host-wise (Araneae), rather than the natural genome size range of the entire order *“Quisvirales”*. For comparison, genome sizes of the multi-family *Picornavirales* and *Nidovirales* vary from two to almost five-fold. It is particularly interesting to find out whether quisviruses with even larger genome sizes exist, given that these viruses don’t share a proof-reading exoribonuclease (ExoN) with the 19-62 kb nidoviruses. Moreover, an increased sampling may verify or expand the current host range of quisviruses that is limited to different species of spiders. Conspicuously, metatranscriptomes of a much larger spectrum of invertebrates were found quisvirus-negative while nidovirus-positive ^23^. This observation indicates that the observed virus-host link for quisviruses and Araneae may be biologically sound.

Interestingly, quisviruses and a provisional family-like group of spider-infecting nidoviruses uniquely share a pair of adjacent REXA and AlkB domains that are encoded in proximity to the RdRp in the viral genome (**Table S5**) ^23^. Both domains may affect RNA replication and/or transcription. AlkB may contribute to genomic repair by demethylation of oxidized nucleotides ^66^. REXA remains an uncharacterized putative ATPase that is genetically segregated in variable locations with the RdRp of many invertebrate nidoviruses and flavi-like viruses ^23,67^.

Also, REXA and AlkB flank the novel AAA+ ATPase domain in pp1ab of quisviruses, and this segregation may be functionally important. In nidoviruses, this AAA+ ATPase locus is occupied by a SF1 helicase. We did not observe top hits to helicases of SF1-SF6 for quisvirus ATPase or another domain, indicating that quisviruses differ from other ssRNA+ viruses with genomes larger than 7.1 kb. Like multimeric ring helicases of SF3-SF6, the quisvirus ATPase appears to adopt a single AAA+/RecA-like fold and shares four key sequence motifs with these helicases (**Figure 3**). In the pair-wise sequence distance space, this ATPase have diverged from the canonical helicase superfamilies as much as they have from each other (**Supplementary Figure 6B**). Collectively, our results are consistent with this ATPase being a helicase, possibly of a new superfamily SF7 comparable to SF3-SF6.

The *“Quisvirales”* is only the second RNA virus order after the *Picornavirales* that encodes a single-AAA+/RecA-like helicase, and its discovery revises the known upper size of the respective virus RNA genomes (**Table S8**). Strikingly, this putative quisvirus helicase and SF3 helicases of the *Picornavirales* differ mostly in sizes of sequence spacers separating four key motifs rather than in the motifs. Compared to the *Picornavirales* counterpart, the putative quisvirus helicase is encoded in the unusual genomic position downstream of the RdRp (as in nidoviruses), which may have affected its role in genome replication, while its expression may be attenuated by the upstream −1 PRF. The available results are compatible with a singular acquisition of a single-AAA+/RecA-like helicase by an ancestral 3CLpro-RdRp encoding virus either upstream or downstream of the RdRp. During subsequent macroevolution, this helicase might have been relocated to the other genomic locus in virus progeny, and the ancestral and descendant helicases gave rise to SF3 and SF7 of the two ssRNA+ virus orders *Picornavirales* and *“Quisvirales”*. Other evolutionary scenarios, including independent acquisitions of helicases in ancestors of the two orders, may be envisioned as well.

While quisviruses encoding a novel SF7 helicase seems the most plausible scenario, these viruses might use the AAA+/RecA-like domain for a function other than nucleic acid unwinding during replication. That would place quisviruses apart from all other known ssRNA+ viruses. Regardless of the actual function of the novel ATPase, the discovery of quisviruses offers unexpected insights into the linkage of RNA virus genome size and helicases and informs studies of orders *Picornavirales* and *Nidovirales*. The quisvirus genomic architecture and proteome implies that large replicases of nidovirus-like complexity evolved repeatedly from an ancestral 3CLpro–RdRp-encoding RNA virus lineage in the phylum *Pisuviricota*.

## Acknowledgements

We thank all colleagues in the scientific community who make their sequence data publicly accessible. We acknowledge the NCBI for providing an elaborate platform to exchange sequencing data. We gratefully acknowledge the computing time made available to us on the high-performance computers Romeo and Barnard at the NHR (Nationales Hochleistungsrechnen an Hochschulen) Center NHR@TUD of the University of Technology Dresden.

## Author contributions

A.E.G. (Conceptualization, Formal Analysis, Supervision, Writing – original draft), D.V.S. (Formal analysis, Software, Writing – review & editing), C.L. (Conceptualization, Formal analysis, Funding acquisition, Software, Supervision, Visualization, Writing – original draft).

## Supplementary data

**Supplementary Figure 1**. Quisvirus multiple nucleotide sequence alignment of the 5’-terminal region of ORF1b encoding the −1 programmed ribosomal frameshifting element as predicted using KnotInFrame. The black bar shows the slippery sequence and its conservation across six quisviruses. The predicted pseudoknot structure is shown for Strandella quadrimaculata quisvirus below the alignment.

**Supplementary Figure 2**. Sequencing read coverage depth of six quisvirus genome assemblies.

**Supplementary Figure 3**. Quisvirus multiple amino acid sequence alignments of PLpro (A) and 3CLpro (B) domains.

**Supplementary Figure 4**. Quisvirus multiple amino acid sequence alignments of RdRp (A), REXA (B), ATPase (C) and AlkB (D) domains.

**Supplementary Figure 5**. Quisvirus multiple amino acid sequence alignments of conserved regions in quisvirus ORF3 and ORF4 products. ORF3 and ORF4 code for, respectively, an immunoglobulin-like (A) and two glycoprotein (GP)-like domains GP1-like (B) and GP2-like (C), respectively (see main text).

**Supplementary Figure 6**. (A) Superimposition of 10 top-ranking quisvirus ATPase models from two independent AlphaFold3 runs; blue: five models from first run, yellow: five models from second run. (B) Network of helicase superfamilies. Nodes represent the quisvirus ATPase or representatives of helicase superfamilies SF1-SF6 of RNA viruses (RNAvir), *Caliciviridae* and calici-like viruses (Calici), *Picornaviridae* and picorna-like viruses (Picorna), DNA viruses (DNAvir) or cellular organisms (see Methods for further details). The widths of the connecting edges are proportional to the maximum sum of posterior probabilities (MSPP) of hits obtained in pairwise HMM-vs-HMM comparisons using HHalign (see Methods for details and Table S3). The comparison of *“*SF3 Calici*”* vs. *“*SF3 Picorna*”* serves as a positive control.

**Supplementary Table 1**. Quisviruses discovered in this study.

**Supplementary Table 2**. Pisuviricota representatives included in phylogenetic analysis.

**Supplementary Table 3**. Results from pairwise HMM-vs-HMM comparisons using HHalign. Values in the sumprobs_max column have been used to calculate weights in helicase network analysis. HHalign was run in (default) local mode and hits may therefore be limited to strongly conserved motifs such as Walker A and B.

**Supplementary Table 4**. Coordinates and identified encoded protein domains of predicted open reading frames of six quisviruses.

**Supplementary Table 5**. Top pHMM-based search hits of quisvirus protein domains against Pfam-A, PDB70, SCOPE70, UniProt20 and internal VDB database.

**Supplementary Table 6**. HHsearch-based profile-vs-sequence search results for predicted (poly)proteins encoded by quisviruses using quisvirus ATPase and SF1-SF6 helicase profiles as query. SF hits within the genomic range of the quisvirus ATPase are in black, otherwise they are in grey. All predicted ORFs were used as target but hits were only observed for ORF1b products. No hits were obtained for SF2, SF5, and SF6.

**Supplementary Table 7**. All pHMM-based search hits of quisvirus ATPase against Pfam-A, PDB70, SCOPE70, and UniProt20. Yellow highlighted are outstanding hits detailed in the text.

**Supplementary Table 8**. Single- and double-RecA-like helicase superfamilies in *Pisuviricota*, location relative to RdRp and relation to genome size.

## Data availability

Transcriptomic data used in this study are publicly available through the SRA and TSA databases at the NCBI. Data accompanying the study have been uploaded to FigShare: DOI 10.6084/m9.figshare.31119181. Viral genome sequences assembled in this study will also be made available through the NCBI Genbank database upon publication. The Virushunter and Virusgatherer tools are available on GitHub: https://github.com/lauberlab/VirusHunterGatherer.

