## Supplementary figures and images for "A large replicase of nidovirus-like complexity in a putative RNA virus order *“Quisvirales”* expands the known diversity of helicase superfamilies"

### Supplementary Figure 1

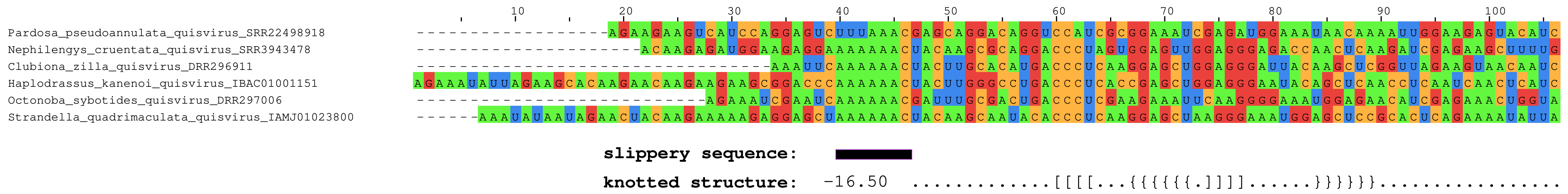

### Supplementary Figure 2

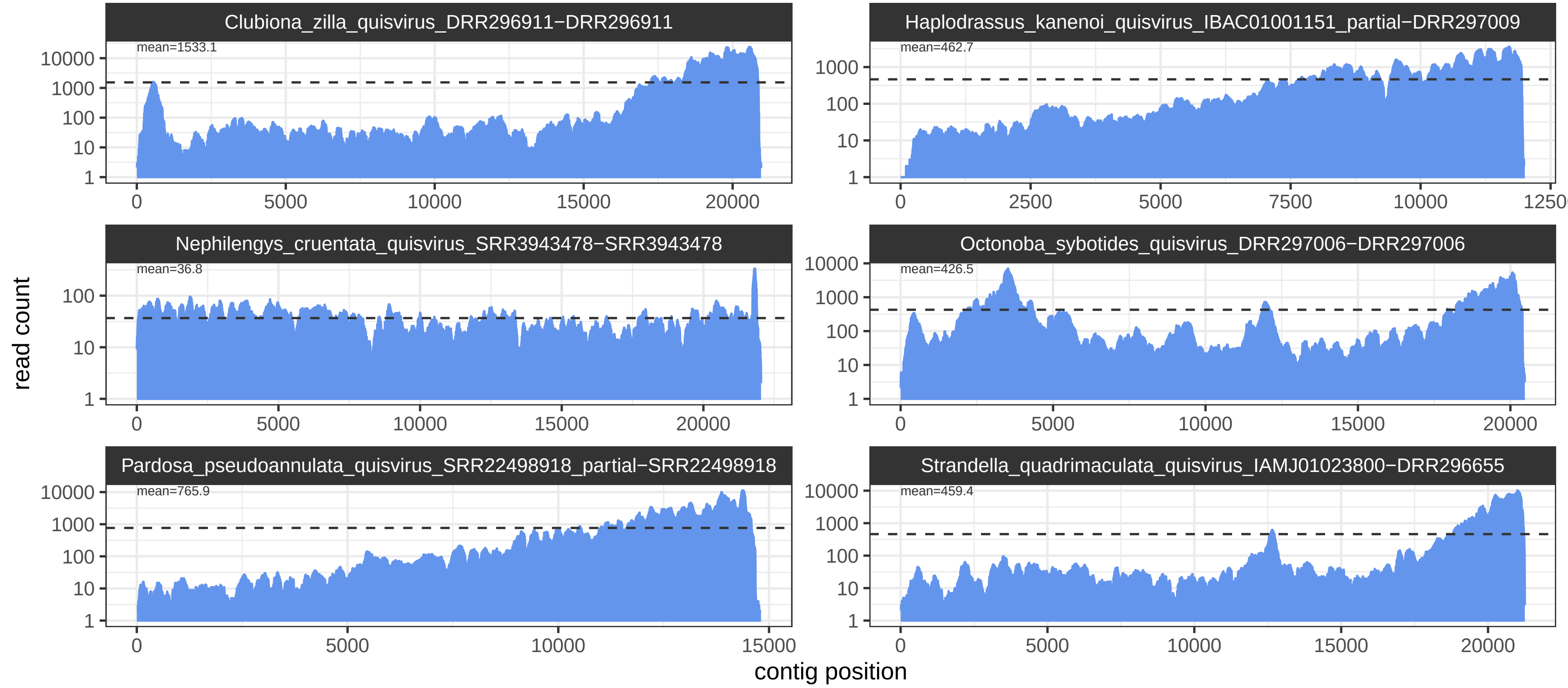

### Supplementary Figure 3

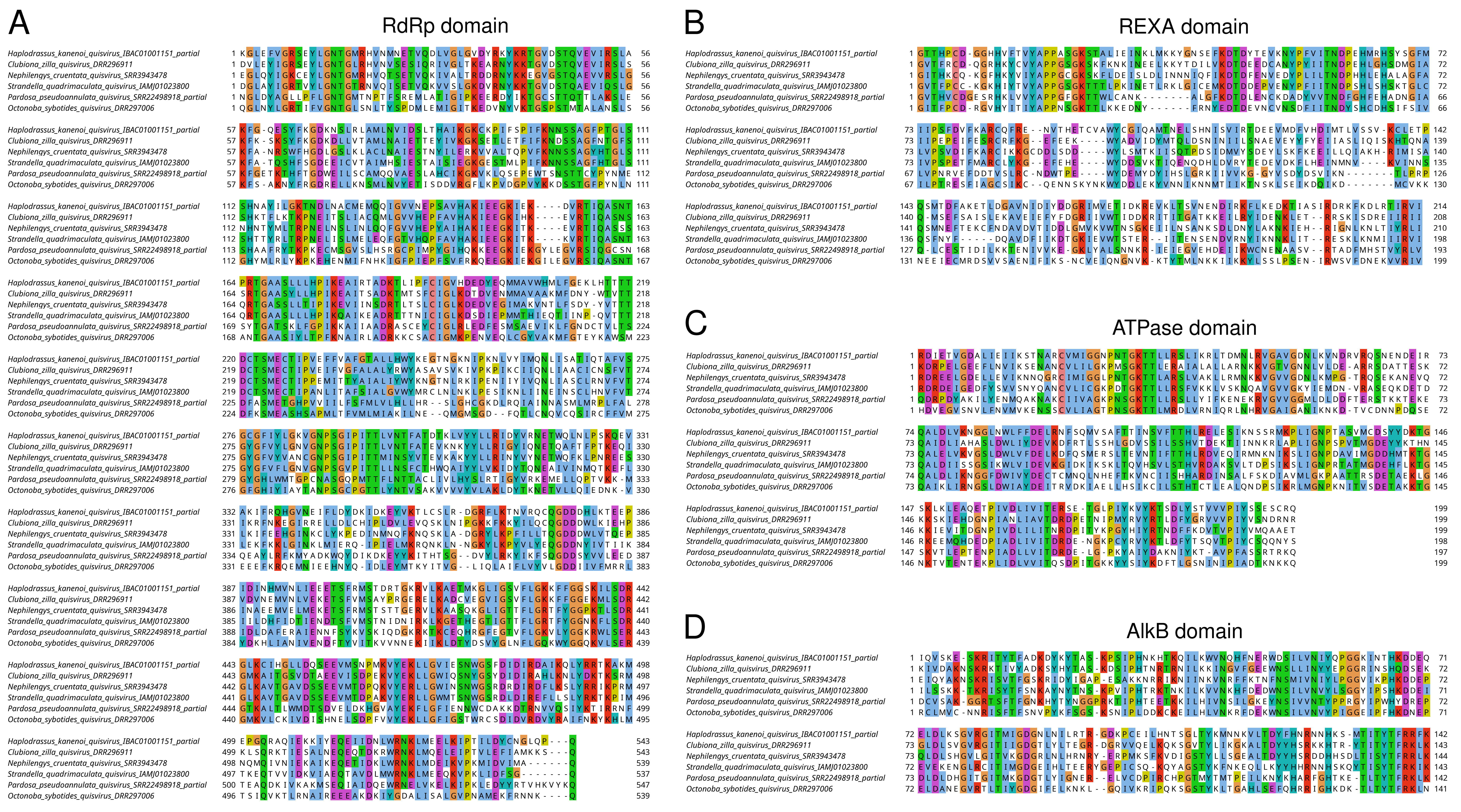

### Supplementary Figure 4

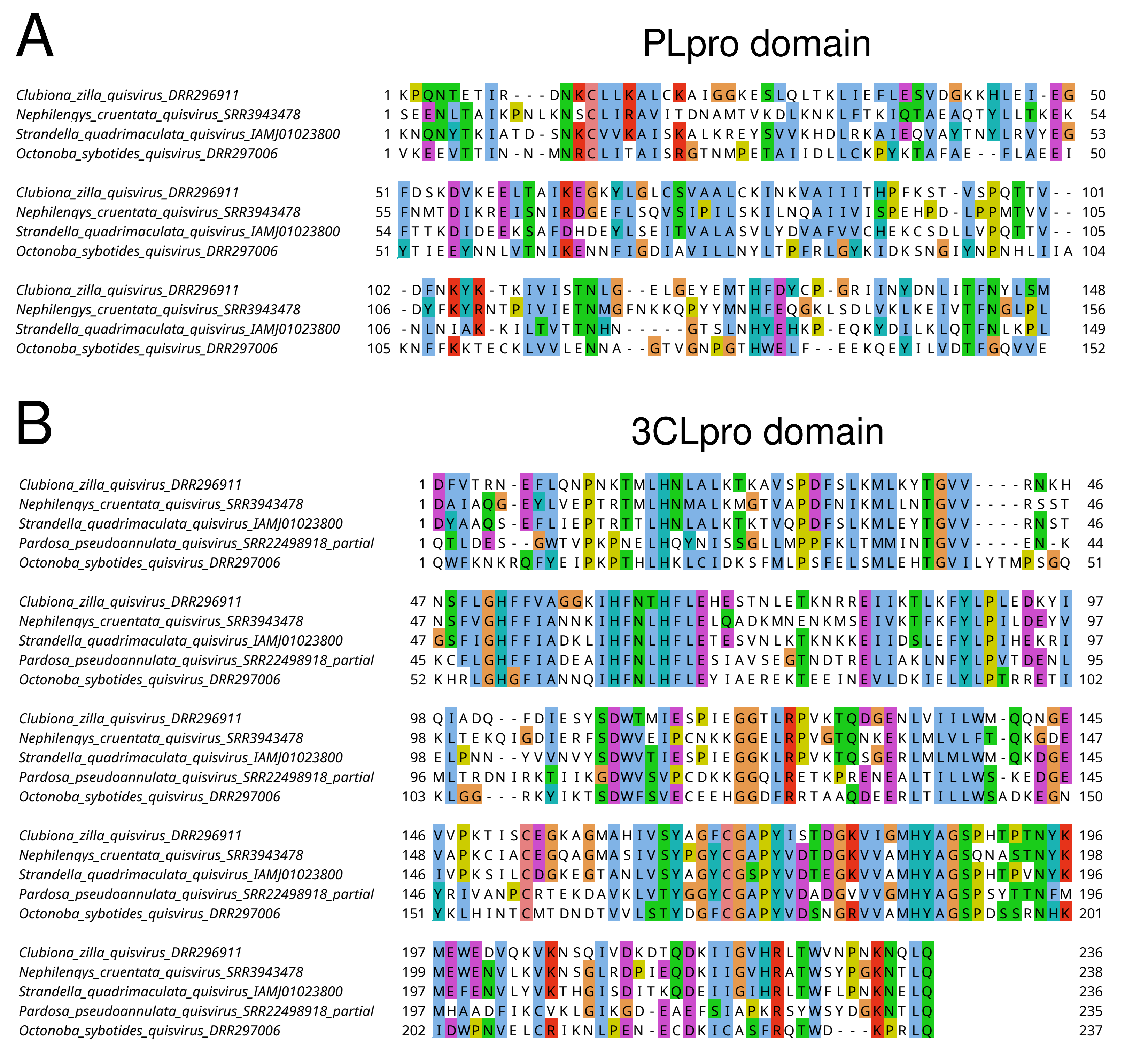

### Supplementary Figure 5

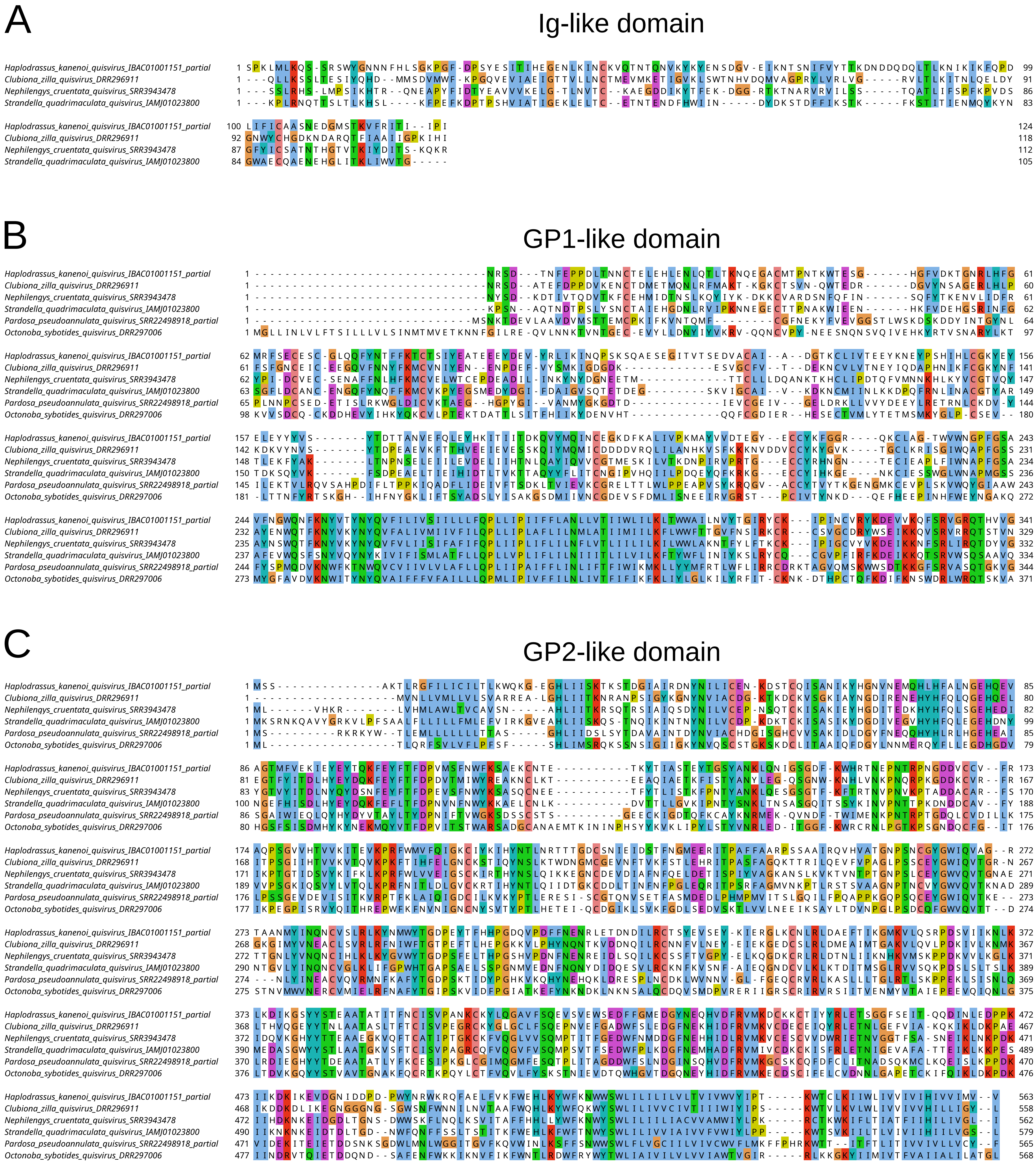

### Supplementary Figure 6

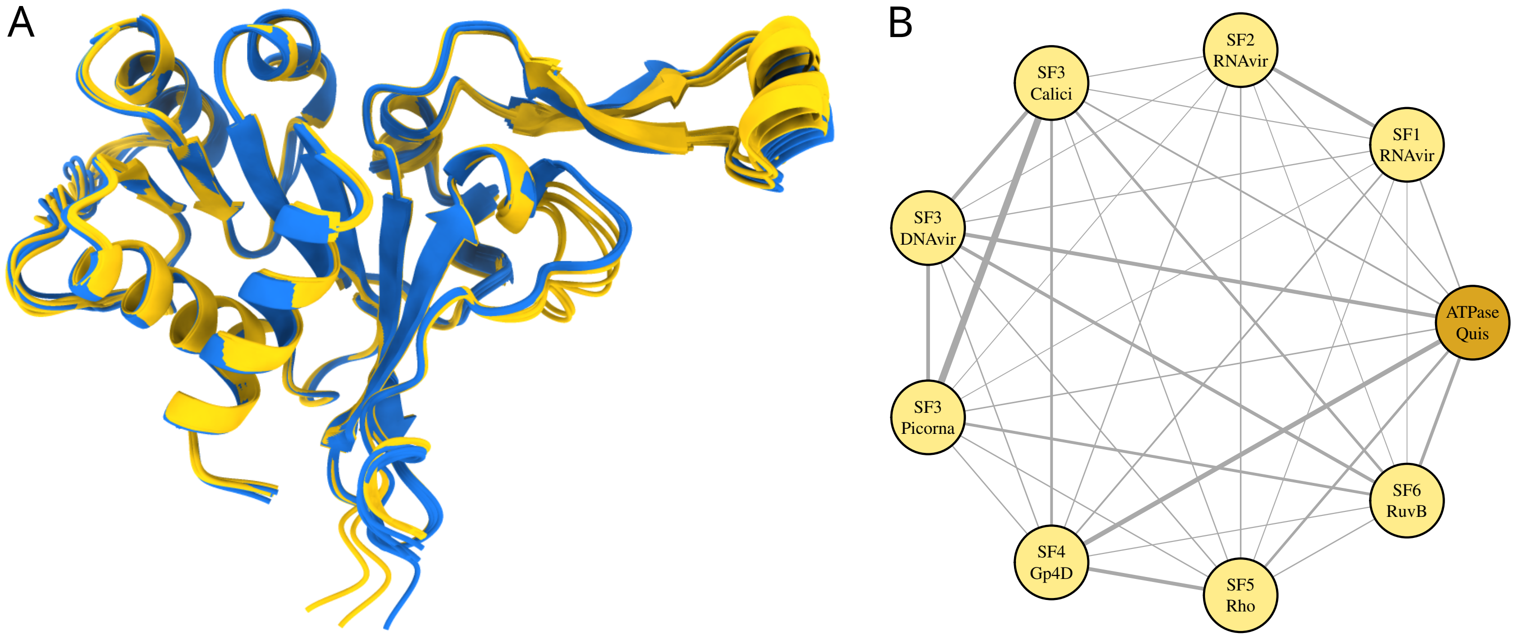
